# Bacterium-Enabled Transient Gene Activation by Artificial Transcription Factor for Resolving Gene Regulation in Maize

**DOI:** 10.1101/2021.02.05.429970

**Authors:** Mingxia Zhao, Zhao Peng, Yang Qin, Ling Zhang, Bin Tian, Yueying Chen, Yan Liu, Guifang Lin, Huakun Zheng, Cheng He, Kaiwen Lv, Harold N. Trick, Yunjun Liu, Myeong-Je Cho, Sunghun Park, Hairong Wei, Jun Zheng, Frank F. White, Sanzhen Liu

**Affiliations:** Department of Plant Pathology, Kansas State University, Manhattan, KS 66506-5502, USA; Department of Plant Pathology, University of Florida, Gainesville, FL 32611-0680, USA; Institute of Crop Sciences, Chinese Academy of Agricultural Sciences, Beijing 100081, P.R. China; College of Forest Resources and Environmental Science, Michigan Technological University, Houghton, MI 49931, USA; State Key Laboratory of Tree Genetics and Breeding, Northeast Forestry University, Heilongjiang Harbin 150040, P. R. China; Innovative Genomics Institute, University of California, Berkeley, CA 94704, USA; Department of Horticulture and Natural Resources, Kansas State University, Manhattan, KS 66506-5502, USA

**Author notes:** These authors contributed equally to this work (Mingxia Zhao, Zhao Peng, Yang Qin). Seeds Research, Syngenta Crop Protection, LLC, Research Triangle Park, North Carolina 27703. DeltaMed Solutions, Inc., 220 Davidson Avenue, Suite 201, Somerset, NJ 08873. National Engineering Research Center of JUNCAO Technology, College of Life Science, Fujian Agriculture and Forestry University, Fuzhou 350002, China (Huakun Zheng).

**Keywords:** transient activation, Xanthomonas, TALe, cuticular wax, maize

## Abstract

Cellular functions are diversified through intricate transcription regulations, and an understanding gene regulation networks is essential to elucidating many developmental processes and environmental responses. Here, we employed the Transcriptional-Activator Like effectors (TALes), which represent a family of transcription factors that are synthesized by members of the γ-proteobacterium genus *Xanthomonas* and secreted to host cells for activation of targeted host genes. Through delivery by the maize pathogen, *Xanthomonas vasicola* pv. *vasculorum*, designer TALes (dTALes), which are synthetic TALes, were used to induce the expression of the maize gene *glossy3* (*gl3*), a MYB transcription factor gene involved in the cuticular wax biosynthesis. RNA-Seq analysis of leaf samples identified 146 *gl3* downstream genes. Eight of the nine known genes known to be involved in the cuticular wax biosynthesis were up-regulated by at least one dTALe. A top-down Gaussian graphical model predicted that 68 *gl3* downstream genes were directly regulated by GL3. A chemically induced mutant of the gene Zm00001d017418 from the *gl3* downstream gene, encoding aldehyde dehydrogenase, exhibited a typical glossy leaf phenotype and reduced epicuticular waxes. The bacterial protein delivery of artificial transcription factors, dTALes, proved to be a straightforward and powerful approach for the revelation of gene regulation in plants.

## INTRODUCTION

Transcriptional regulation is essential for cellular differentiation and responses to environmental signals. Transcription factors (TFs) are key components for modulating gene expression and an understanding TF function is fundamental to elucidating gene regulation networks. Traditional approaches to transcription pathway analysis involve ectopic expression, in some cases transiently, and genetic mutation. Further analyses involve the analysis of TF binding sites, including chromatin immunoprecipitation sequencing (ChIP-Seq) and the *in vitro* DNA affinity purification sequencing (DAP-Seq) (Bartlett et al., 2017). All of the approaches have advantages and limitations (Lai et al., 2019). Ectopic expression or knockouts are typically constructed to understand phenotypical and transcriptional consequences of a particular TF, and, at the same time, are not readily available or require considerable time to construct. Genome-wide transcriptional changes through the transient gene activation may offer another approach (Gleba et al., 2014). However, transient expression methods can be limiting, depending on species. Emerging nanomaterial technologies offer a potential option for delivery of nucleotides and proteins, and the techniques efficiently overcoming the barrier from plant cell walls are still evolving (Cunningham et al., 2018; Demirer et al., 2020).

Bacteria, principally pathogenic species, although not all, have evolved secretion systems to inject proteins into host cells to induce changes in the host metabolism, and facilitate colonization of host tissues (Costa et al., 2015; Deslandes and Rivas, 2012; Block et al., 2008). The type III secretion system (T3SS) is such a supramolecular complex that delivers bacterial proteins (effectors) to target cells (Green and Mecsas, 2016). Many plant pathogens of the genus *Xanthomonas* require a functioning T3SS for virulence and cause diseases on hundreds of plant species, including most major crop species (White et al., 2009; Büttner and Bonas, 2010). The Transcriptional-Activator Like effector (TALe) family is a group of type III effectors that have diverse functions for host cell manipulations, with the primary function of directing expression of specific disease susceptibility genes. The N-terminus of a TALe contains the T3SS secretion signal and the C-terminus processes domains for eukaryotic nuclear localization and transcription activation (Yang et al., 2000; Zhu et al., 1998; Van den Ackerveken et al., 1996). The central repetitive sequence consists of a variable number of repeats, each of which contains nearly identical 34-35 amino acid residues and variable di-residues (RVD) at the 12^th^ and 13^th^. The RVD of a repeat determines the specific recognition of a nucleotide base of four DNA nucleotides (Boch et al., 2009; Moscou and Bogdanove, 2009). The revelation of specific recognition between RVD and nucleotide bases provides a rationale for the construction of artificial, or designer, TAL effectors (dTALes) to target specific DNA sequences (Joung and Sander, 2013; Li et al., 2013b).

Here, we used TALe-mediated targeting activation of transcription as delivered by the maize pathogen *Xanthomonas vasicola* pv. *vascularum* (Xvv) (Perez-Quintero et al., 2020) to characterize the regulation of cuticular wax development. Cuticular waxes are derivatives of very-long-chain fatty acids (VLCFAs), which are produced through cyclic reactions that add two carbons per cycle (Kunst and Samuels, 2003; Lee and Suh, 2013). Cuticular waxes are secreted through the plasma membrane to the plant surface, providing a hydrophobic barrier to protect plants from water loss and other environmental stresses (Fehling and Mukherjee, 1991). In maize, mutants with reduced cuticular waxes can hold water droplets on leaves with water spraying, which is referred to as the glossy phenotype. To date, more than 30 glossy loci have been discovered by mutants, and at least 11 glossy genes were found to be responsible to glossy leaf phenotype, including *gl1*, *gl2, gl3, gl4, gl6, gl8, gl13, gl14, gl15, gl26*, and *cer8* (Tacke et al., 1995; Li et al., 2019, 2013a; Liu et al., 2012; Moose and Sisco, 1996; Zheng et al., 2019; Hansen et al., 1997; Xu et al., 1997; Liu et al., 2009). Among them, the *gl13* gene encodes an ABC transporter functioning in secretion of cuticular waxes through the plasma membrane (Li et al., 2013a). The *gl15* gene encodes an AP2-like TF, which does not directly participate in the biosynthesis of cuticular wax but functions in juvenile-to-adult transition of epidermal cells (Moose and Sisco, 1996). In this study, the artificial transcription factors, or dTALes, were used to activate *gl3*, an apparent early TF gene in the biosynthesis of cuticular waxes, and identify the downstream genes of GL3 and genes related to the cuticular wax pathway.

## RESULTS

### A bacterium-enabled protein delivery system in maize

The Xvv strain Xv1601, a pathogen of maize, was used for delivery, and, based on previous sequence analysis, is free of TALe genes (Perez-Quintero et al., 2020). Phylogenetic analysis of 10 *Xanthomonas* species indicated that Xvv is genetically close to *Xanthomonas oryzae* (Xo) (**Supplemental Figure 1**). Xv1601 does contain a T3SS gene cluster that is syntenic with the clustered genes from a reference Xo strain PXO99^A^, which is known to deliver TALes during infection (**Figure 1A**). A knockout mutant of *hrcC* (*hrcC^-^*), an essential gene of the T3SS, dramatically reduced the virulence of Xv1601, indicating T3SS is critical for the bacterial virulence (**Figure 1B**). To test the ability of Xv1601 to deliver proteins to intact maize leaf cells, a plasmid bearing a green enhanced fluorescent protein (eGFP) gene fused to the type III secretion signal of AvrBs2 was constructed (**Supplemental Figure 2**) (Minsavage, 1990). To enhance detection, a nuclear localization signal was incorporated into the protein (Khang et al., 2010). Upon inoculation of leaf tissue, GFP signals were detected in host cells (**Figure 1C**). In this case, GFP was localized to the nucleus.

**Figure 1.**
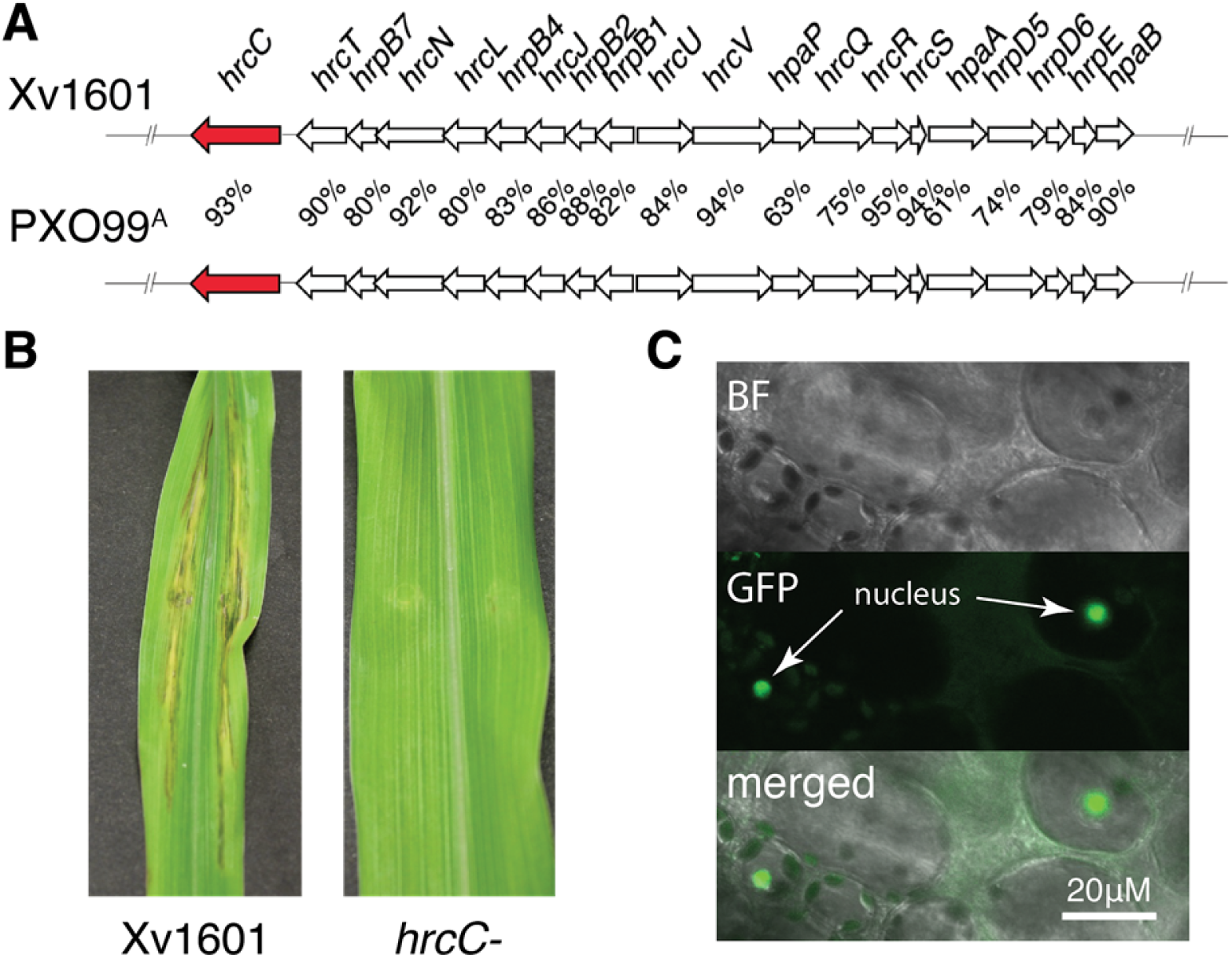
The T3SS of Xvv functions delivering proteins to intact cells. (**A**) Comparison of the type III secretion gene cluster of Xv1601 and the *Xanthomonas oryzae* strain PX099^A^. The DNA identity between each orthologous pair is listed. The *hrcC* genes are highlighted in red. (**B**) Leaf phenotype five days after inoculation with the wildtype strain (Xv1601) and the *hrcC* knockout mutant strain. (**C**) Bright field (BF) and fluorescence (GFP) images of maize cells after 24 h of the infection with bacteria carrying a gene of AvrBs2::T3SS signal peptide-NLS::eGFP-NLS (**Supplemental Figure 2**). T3SS: The type III secretion system.

### TALe-induced expression of a host gene

As Xv1601 does not contain any endogenous TALe genes, we tested the ability of the strain to deliver dTALes based on the ability to induce a targeted host gene. Two dTALes, referred to as dT1 and dT2, were constructed to target two non-overlapping 16-bp regions of the *gl3* promoter (**Figure 2A**, **2B**). Both dT1 and dT2 targeted regions are close to two predicted TATA boxes, which are 5 bp and 48 bp upstream of the transcription start site, respectively. Expression of *gl3* was observed in seedling leaves.

**Figure 2.**
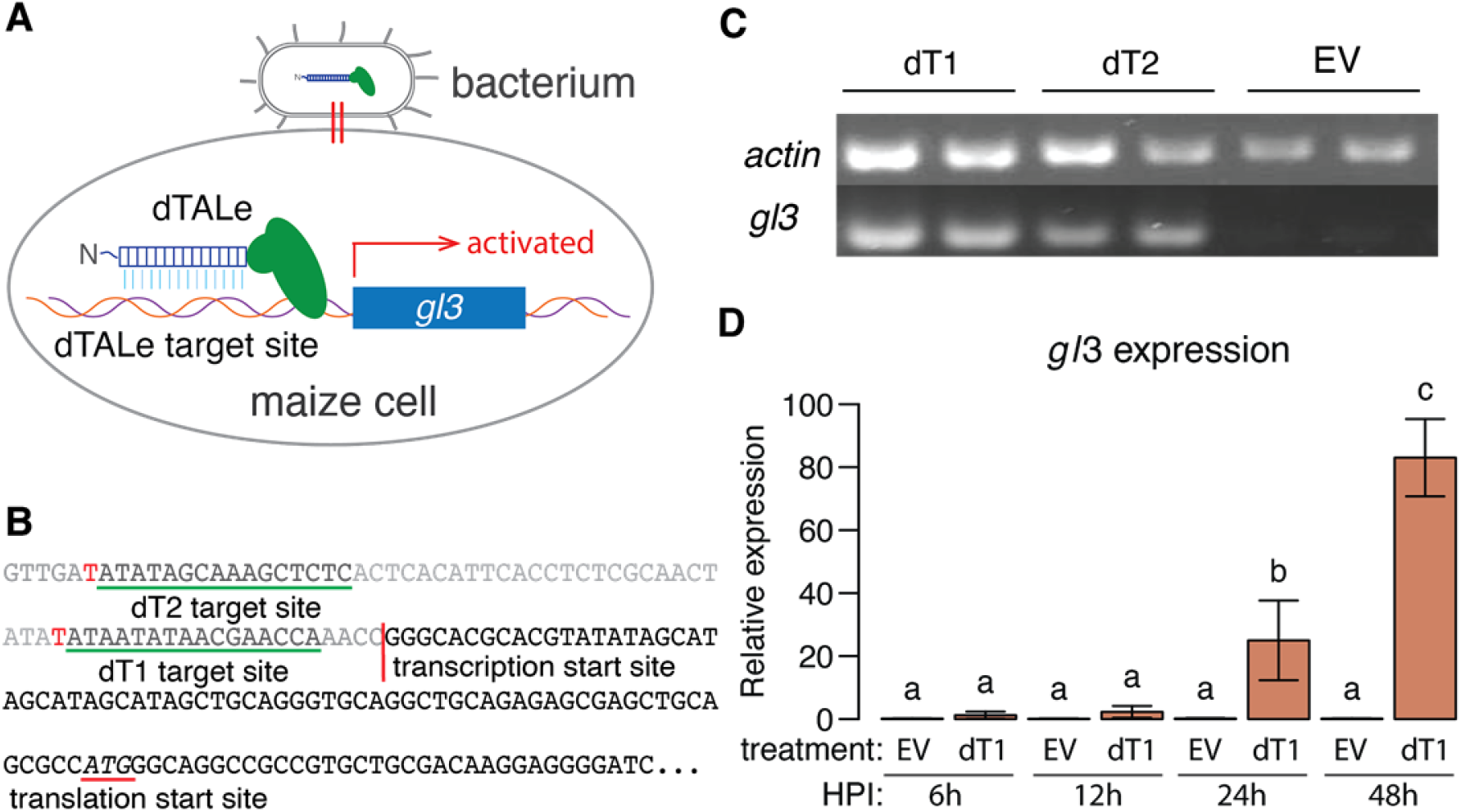
dTALe-dependent *gl3* gene expression. (**A**) Schematic of bacterium-mediated delivery of dTALes for the expression activation of maize *gl3*. (**B**) The target sequences for dT1 and dT2 (underlined in green). The transcription start site is indicated by a vertical red line. The translation start site ATG for GL3 is underlined in red. (**C**) Semi-RT-PCR of the *gl3* expression from 14-day old seedling leaves. Treatments with two replicates are shown for bacteria carrying either dT1, dT2, or the empty vector (EV). The constitutively expressed *actin* gene was used for loading controls. (**D**) qRT-PCR of the *gl3* expression at 6, 12, 24, and 48 hours post inoculation (HPI). The bar heights are the average of three biological replicates per treatment per time points. Error bars represent standard deviation. Values with the same letter do not differ at the significance level of 0.05 as determined by ANOVA and Tukey’s honestly significant difference.

However, expression dropped to the undetectable levels by 14 days after planting (**Supplemental Figure 3**). Therefore, 14-day seedlings were used to test for dTALes-mediated induction of *gl3*. Bacterial strains carrying either dT1 or dT2 activated *gl3* expression by 24 h after the bacterial inoculation (**Figure 2C**). Compared with dT2, dT1 promoted stronger induction of *gl3* as measured by quantitative reverse transcription PCR (qRT-PCR) (**Supplemental Figure 4**). A time-series analysis of the *gl3* expression due to dT1 showed that relative expression reached 22 fold and 82 fold at 24 h and 48 h after inoculation, respectively, compared to the empty vector (**Figure 2D**).

### GL3 downstream genes identified through RNA-Seq

To determine the genes regulated by GL3, RNA-Seq was performed using tissues after treatments with bacteria carrying dT1, dT2, and the empty vector (EV). The basal expression level of *gl3* in young leaves was low, while treatments with dT1 or dT2 exhibited 191 and 74 fold induction of *gl3*, respectively (**Figure 3A, Table 1**). The comparison of dT1 with the EV control identified 1,249 differentially expressed genes (DEGs) at the false discovery rate (FDR) of 5%, among which 499 were up-regulated. A comparison of dT2 versus EV resulted in 430 DEGs were identified at the FDR of 10%, of which 156 were up-regulated (**Figure 3B**). Note that a higher FDR value used in dT2 is due to a lower level induction of the *gl3* expression. The 92 common up-regulated DEGs of dT1 and dT2 and 54 common down-regulated DEGs were deemed as the *gl3* downstream genes (excluding *gl3*). Gene ontology (GO) analysis showed that the genes related to fatty acid biosynthesis and the endoplasmic reticulum (ER) are overrepresented in the 92 up-regulated genes (**Figure 3C**).

**Figure 3.**
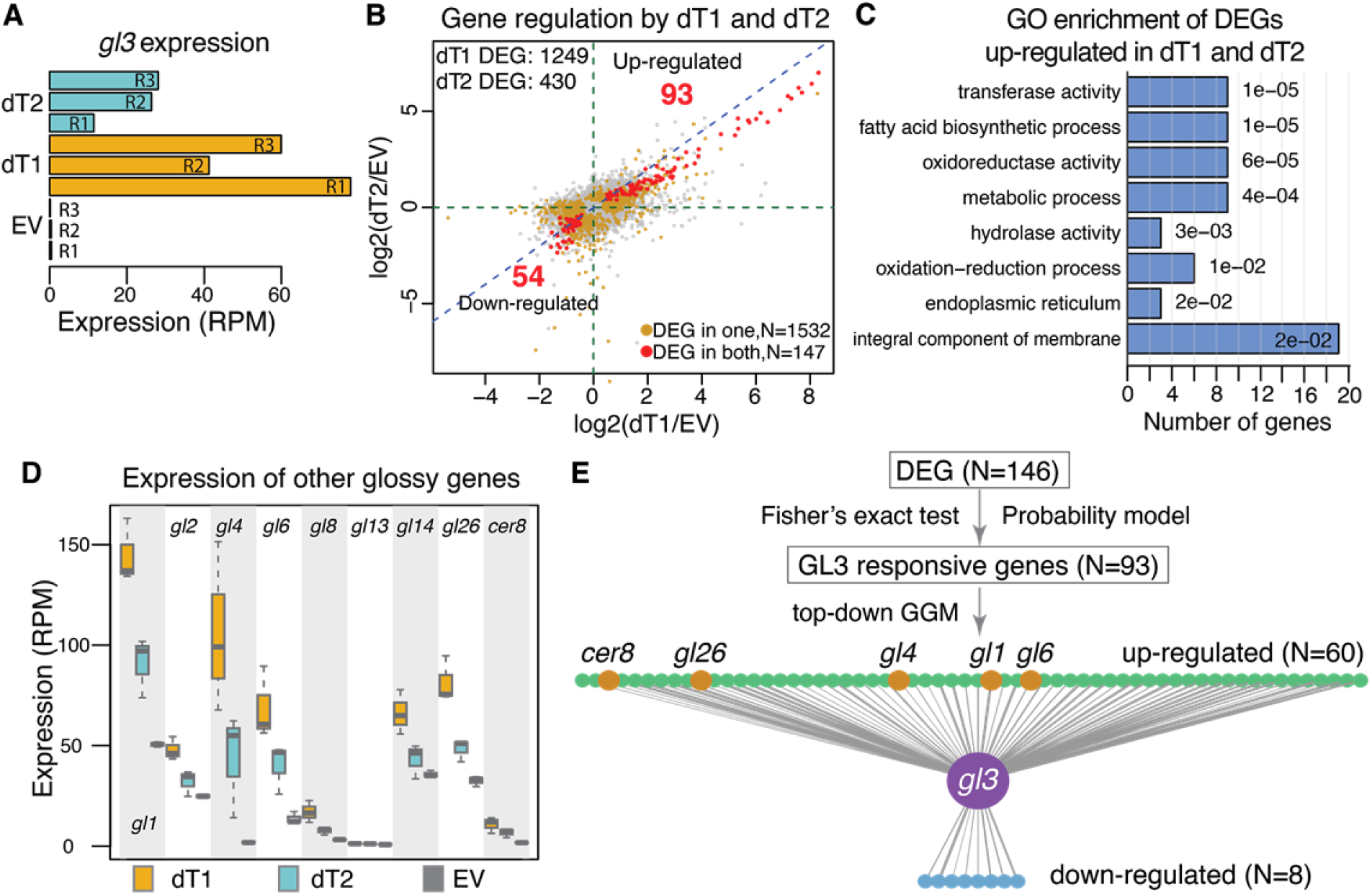
Gene expression associated with TALe-dependent expression of *gl3*. (**A**) Expression in RPM (reads per million reads) of *gl3* from RNA-Seq data. R1-R3 represent biological replicates. The treatment groups EV, dT1, dT2 stand for constructs of empty vector, dT1, and dT2, respectively. (**B**) Scatter plot between log2 fold changes of gene expression in the comparison of dT1 versus EV and that in the comparison of dT2 versus EV. The 93 genes up-regulated by both dT1 and dT2 include *gl3*. Gray, orange, and red points represent unaffected, DEGs in one comparison, and DEGs in both comparisons, respectively. (**C**) Gene ontology (GO) enriched in the DEGs in both comparisons. Numbers besides bars are p-values of GO enrichment tests. (**D**) Expression in RPM of nine glossy genes that affect cuticular wax accumulation in three treatment groups. (**E**) Direct regulation by GL3 indicated by the top-down GGM analysis. The upper layer listed up-regulated genes by *gl3* and the bottom layer listed down-regulated genes. The thickness of connection lines represents the number of interferences by GL3 for each gene. Glossy genes are highlighted in orange.

**Table 1.**
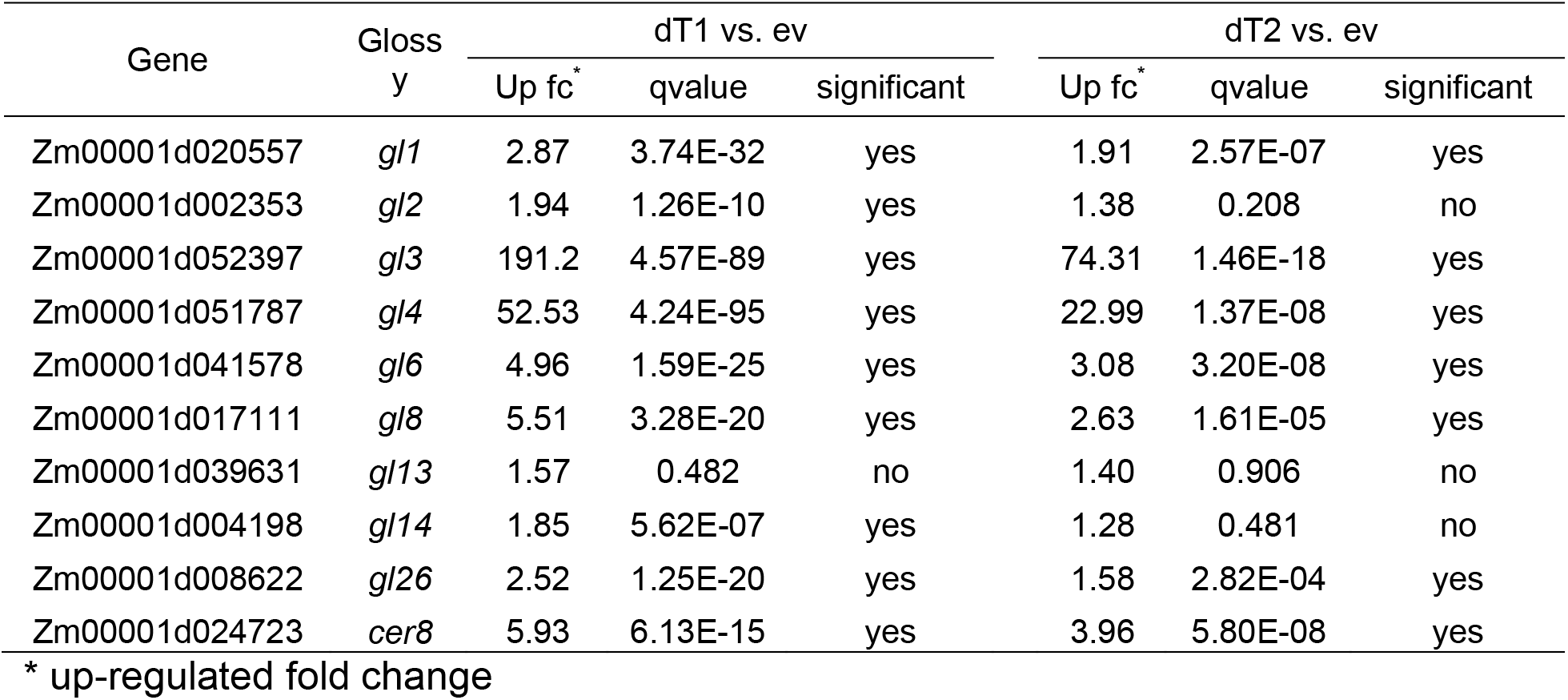
Differential expression of known glossy genes

Of nine known glossy genes, which do not include *gl3* or *gl15*, six were among the 92 up-regulated DEGs that were up-regulated by both dTALes, and additional two genes were only up-regulated by dT1 (**Table 1**). All eight genes showed the same regulation pattern by two dTALes, of which dT1 exhibited a stronger induction (**Figure 3D**). The only glossy gene that was unaffected by the *gl3* induction is *gl13*, which is an ABC transporter functioning in the secretion of cuticular waxes across the plasma membrane (Li et al., 2013a). Besides the known glossy genes, 86 additional genes were up-regulated by the dTALes including six genes encoding 3-ketoacyl-CoA synthases (KCS) as *gl4* does (Liu et al., 2009), two genes encoding HXXXD-type acyltransferase related proteins similar to *gl2* (Tacke et al., 1995), three genes encoding GDSL esterase/lipase proteins, which was reported to be involved in wax biosynthesis (Tang et al., 2020), and two genes encoding aldehyde dehydrogenases (**Supplemental Data Set 1**). The 54 down-regulated genes were identified in both dT1 and dT2 comparisons with the EV group, which do not include any known glossy genes. Most glossy genes were previously reported to be clustered in a turquoise module of a gene co-expression network (GCN295) constructed using 295 RNA-Seq data (Zheng et al., 2019). Of the 92 genes up-regulated by *gl3*, 61 are present in GCN295, and 38/61 were assigned to the turquoise module. In contrast, only three genes from the 54 genes down-regulated by dTALes are in the turquoise module. Collectively, from the RNA-Seq results, the *gl3* gene appeared to be a master regulator positively modulating biosynthesis of cuticular waxes.

A conventional neural network (CNN) deep learning approach was used to determine the probability that a gene is regulated by GL3 from 739 publicly available RNA-seq datasets of inbred line B73. To train the prediction model, the gene pairs of TFs and their targeted genes mapped from *Arabidopsis* gene regulation data were used as the positive pairs (Yilmaz et al., 2011), and the random gene pairs that did not overlap with positive pairs were used as the negative control pairs. The deep learning predicted that 59.6% GL3 downstream genes were regulated by GL3 with a probability of at least 0.8, while 17.7% of the 594 control genes that were unaffected by both dT1 and dT2 were predicted (**Supplemental Table 1, Supplemental Data Set 2**). The *in silico* prediction supported that most of the *gl3* downstream genes revealed are regulated by *gl3*.

### Probability-based identification of GL3 directly regulated genes

The GL3 downstream genes are either directly or indirectly regulated by GL3. A top-down Gaussian graphical model (GGM) algorithm was employed to find the genes that were likely to be directly regulated by GL3 (Lin, Li et al. 2013, Wei 2019, Wei, Liu et al. 2020). From the 146 GL3 downstream genes, the algorithm first identified 93 GL3 responsive genes that had high concordance in expression levels with *gl3* expression with the expression data from the dT1, dT2, and EV RNA-seq experiments (**Supplemental Data Set 3**). The expression data of *gl3* and the 93 GL3 responsive genes were then used to infer the directly regulated genes of GL3. Briefly, two genes from the GL3 responsive genes were combined with *gl3* to form a triple gene block for evaluation. If the corrected p-value with multivariate delta method (Methods) for each triple gene block is less than 0.05, *gl3* was scored as to interfere with the two responsive genes once. All triple gene blocks were evaluated, and the interference frequency for each gene was calculated. As a result, 68 genes that were interfered by *gl3* were identified as direct targets of GL3, including 60 up-regulated and 8 down-regulated genes (**Figure 3E**). The remaining 78 genes from 146 GL3 downstream genes are likely to be indirectly regulated by GL3 (**Supplemental Data Set 3**). Five glossy genes that were associated with dT1 and dT2, namely, *gl1*, *gl4*, *gl6*, *gl26*, and *cer8*, were predicted to be directly regulated by GL3, indicative of the direct regulation role of GL3 in biosynthesis of cuticular waxes.

### A *gl3* downstream gene functions in cuticular wax accumulation

Due to the presence of most known glossy genes in the DEGs up-regulated by *gl3*, the dTALe up-regulated genes may contain unknown genes involved in biosynthesis of cuticular waxes. The genes that were up-regulated by both dTALes and assigned to the turquoise module of GCN295 were selected as the candidate glossy genes for the validation. Ethyl methanesulfonate (EMS) induced mutants of four candidate genes were obtained from a maize EMS mutant stock collection (Lu et al., 2018). All mutants were screened for the glossy phenotype. No glossy phenotype was observed for mutants carrying premature stop codons in the three genes Zm00001d046642, Zm00001d028241, and Zm00001d032719, which encode GDSL esterase/lipase, 3-ketoacyl-CoA synthase, and long-chain-alcohol oxidase FAO4B, respectively (**Supplemental Table 2**). Zm00001d017418, which encodes aldehyde dehydrogenase, is up-regulated by treatments with either dT1 or dT2 (**Figure 4A**). The EMS mutant (*ems4-12ff6f*) with a premature stop codon in the second exon of Zm00001d017418 displayed a glossy phenotype, indicating reduced accumulation of cuticular waxes (**Figure 4B, 4C**). Total leaf waxes on *ems4-12ff6f* mutants were reduced by ~40% of that in the wildtype (**Figure 4D**). Microscopic examination of wax components on the leaf surface revealed fewer wax crystals accumulated on leaf surfaces of mutant lines as compared to wildtypes (**Figure 4E**). Wax component analysis found a decrease in C30 and longer chain primary alcohols, alkanes, and fatty acids (**Supplemental Figure 5**).

**Figure 4.**
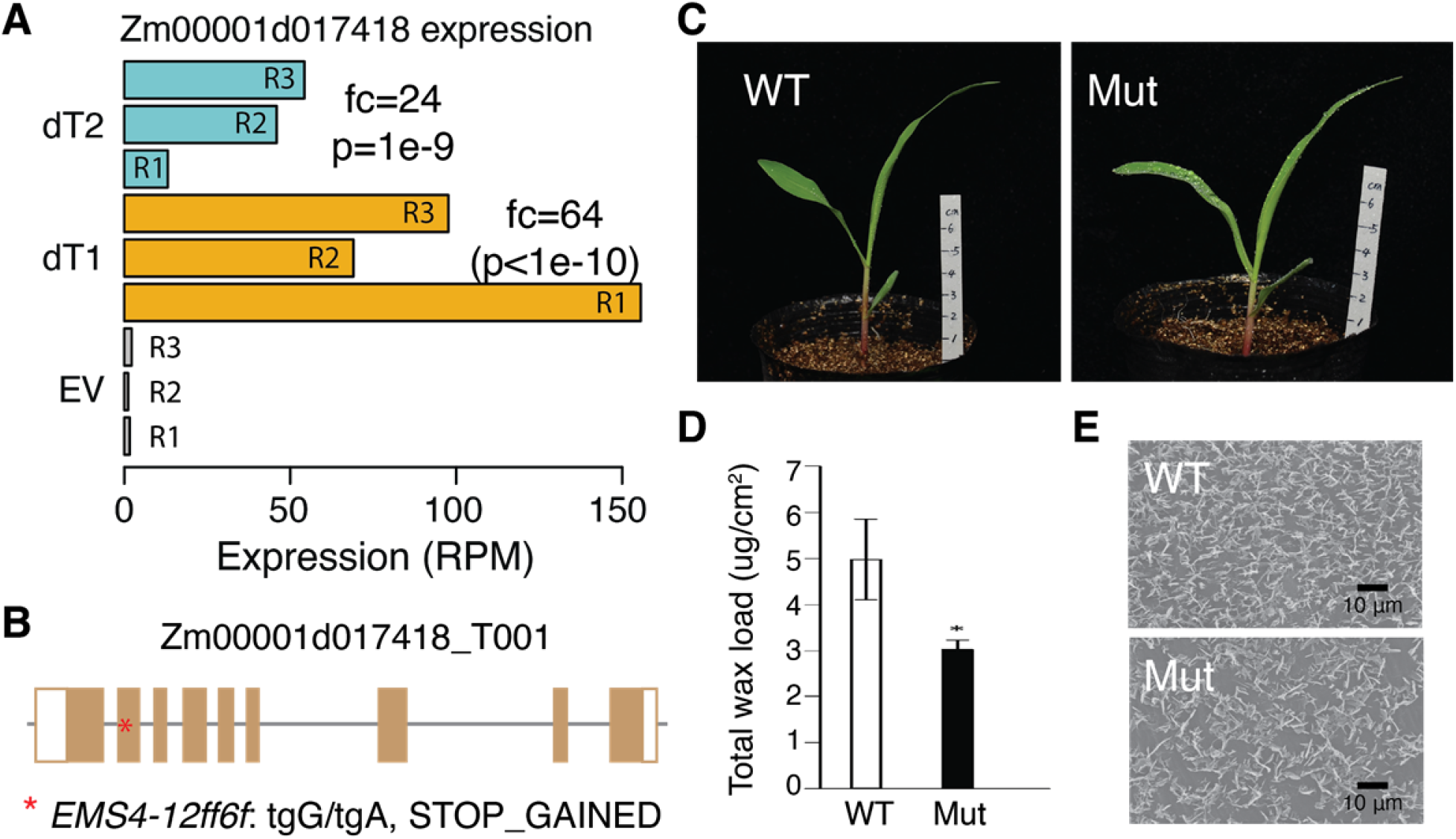
A new glossy gene Zm00001d017418. (**A**) Expression of the candidate gene Zm00001d017418. R1-R3 represent biological replicates. The treatment groups EV, dT1, dT2 stand for constructs of empty vector, dTALe 1, and dTALe 2, respectively. fc, fold change in expression relative to EV; p, adjusted p-value from RNA-Seq analysis. (**B**) Gene structure of the isoform of Zm00001d017418_T001. Boxes are exons and blank boxes represent untranslated regions. Start points at the EMS mutation location, which produces a premature stop codon. (**C**) The visible glossy phenotype of the EMS mutant and the wildtype (WT). Water drops were present on the surfaces of mutant seedling leaves due to reduced epicuticular waxes. (**D**) Total cuticular wax loads and wax components of mutants and wildtypes. (**E**) Epicuticular wax contents on the leaf in the wildtype and the mutant detected via scanning electron microscopy (SEM, x10,000 magnification).

## DISCUSSION

Here, the maize pathogen Xvv and the bacterial T3SS system were used for protein delivery into intact maize cells and, in this specific case, characterization of the cuticular wax pathway. Although considered destructive of plant tissue, *Xanthomonas* species are best considered as hemi-biotrophic in that the pathogens interact with intact cells for some time before destruction of the cells is evident in compatible interactions.

In the initial demonstration, the T3SS signal of the effector AvrBs2 was used to direct GFP to intact cells. NLS was added to the effector to concentrate the protein in nuclei, both as evidence for intact cellular organelle and to facilitate detection of the protein in plant cells. For a demonstration of the utility, the approach was used to study consequences of ectopic expression of the MYB TF GL3 through induction by synthetic TALe, or dTALe, transcription factors. TALe effectors are particularly useful for the approach as TALes already have T3SS secretion signal sequences and NLS for localization into host cell nuclei. Although Xvv does not contain endogenous TALes, the presence of TALes with biological function in disease, including TALes that target host TFs, in closely related strains indicated that TALe delivery would be successful.

Previous experience with the so-called American strains of Xo, which also lack endogenous TALe genes, has indicated that TALes can be delivered by TALe-deficient strains (Tran et al., 2018). In this study, two dTALes were targeted to two separate DNA sequences in the promoter of *gl3*. Both dTALes resulted in *gl3* induction as shown by both qRT-PCR and RNA-Seq. In addition to expression of *gl3*, evidence was obtained that GL3-regulated genes were identified as a consequence of dTALe activation of *gl3*. The best evidence is that one of the apparent GL3 downstream genes, Zm00001d017418, was up-regulated and has the glossy phenotype, which results in reduced wax deposition on leaves, when mutated. The failure of displaying a glossy phenotype of mutants from three other genes does not indicate no involvement in GL3-dependent events and might be due to the functional redundancy in the maize genome. In addition, GL3 downstream genes include most known glossy genes. The results indicate the master regulatory role of GL3 in biosynthesis of cuticular waxes, and provide strong evidence for the efficacy of the dTALe system for revelation of gene regulations. In the future, experimental data can be generated to further examine the binding motif of the TF GL3. Also, given the fact that the *gl3* gene is largely silenced at the adult stages (Zheng et al., 2019), it would be interesting to examine the impacts on the wax biogenesis from the constitutive expression of *gl3*. More genes, particularly TFs downstream of GL3, could be examined through dTALe activation and/or knockouts, to further understand the regulation network of cuticular wax biosynthesis.

The dTALe activation system is easy to manipulate, and *Xanthomonas* strains are easy to culture. Besides the simplicity of the system, it is flexible to control the bacterial load by adjusting the concentration and amount of bacterial inoculum. At the same time, limitations need to be considered for the experimental design. First, multiple independent dTALes are needed to reduce the impacts of off-targeting gene induction. Multiple dTALes help discriminate between off-target gene induction with the idea that independent binding sites will not result in induction of the same off-target genes. Given that no specific domain other than a “T” preceding the binding site is required (Moscou and Bogdanove, 2009), candidate dTALe binding sites are relatively abundant. Second, bacterial infection and other T3SS effectors could potentially interfere with host gene expression or host protein function, if related to defense responses. The bacterium carrying an empty vector as the control, as implemented in this study, should largely reduce the impacts from bacteria.

The downstream genes of a dTALe targeted gene include direct or indirect targets of the dTALe binding gene. Based on expression patterns, direct and indirect regulations are distinguishable with dedicated computational algorithms. The top-down GGM algorithm, with the input of a short time-course data, had been shown to separate the directly from indirectly regulated genes with more than 90% accuracy (Wei, 2019; Lin et al., 2013) and about 80% accuracy for RNA-Seq data from stably transgenics lines (Wei et al., 2020). In this study, no time-course data were generated. However, the variation of *gl3* expression induction within a dTALe treatment, probably due to the variation in the bacterial amount injected during inoculation, and between two dTALes mimics multiple levels of *gl3* induction as in time-course experiments. The data, therefore, enabled the top-down GGM algorithm to identify the genes that closely followed expression changes of *gl3*, which were deemed to be directly regulated genes. Alternatively, the result from dTALe experiments could be combined with the results from DAP-Seq or CHIP-Seq that examines protein-binding sites to identify direct targets.

The *Xanthomonas* bacteria can be used as a general tool for protein delivery to plant host cells. *Xanthomonas* bacterial strains are available for most crops and have a well-documented ability to deliver diverse proteins. This gene activation through dTALes represents a unique system to study transcriptional regulation. The protein delivery system can also be utilized to study plant-pathogen interactions. For example, any effector gene can be engineered to the *Xanthomonas* bacterium and delivered to host cells for examining defense responses. To reduce pathogenic effects from *Xanthomonas*, the bacterium can be modified for the reduced host cell toxicity and a higher capacity for protein delivery.

## METHODS

### Genetic materials

The bacterium Xv1601 is pathogenic on maize (Perez-Quintero et al., 2020). A *hrcC* knockout mutant was generated following protocol previously described (Peng et al., 2016). The maize inbred line A188 (PI 693339) were obtained from the North Central Regional Plant Introduction Station and maintained at Kansas State University. Plants were grown in a growth chamber under 27°C during daytime and 21°C at night with 16 hours of photoperiod. EMS mutants were ordered from the Maize EMS induced Mutant Database (MEMD) (Lu et al., 2018).

### Design and assembly of protein delivery constructs

The pENTR™ 11 Dual Selection Vector (Thermo Fisher Scientific, USA) was digested by KpnI and XhoI, and the DNA fragments containing AvrBs2 promoter, the type III signal peptide, and eGFP were cloned into the digested plasmid according to NEBuilder HiFi DNA Assembly protocol (New England Biolabs, USA). The assembled entry construct was then cloned into the broad host-range vector pHM1 by the Gateway cloning (**Supplemental Figure 2**) and transformed into Xv1601 strain by electroporation using Bio-rad Micropulser (Peng et al., 2019).

### Design and assembly of dTALe

The promoter elements targeted by TAL effectors are typically, not far away, upstream of transcriptional start sites (Moscou and Bogdanove, 2009). Based on previous reports, most TAL effectors (e.g., PthA4, AvrBs3, PthXo2, PthXo3, AvrXa7, PthXo6 and PthXo7) binded at TATA box regions while some (e.g., PthXo1 and Tal8) targeted the regions a few base pairs upstream of TATA boxes (Kay et al., 2007; Sugio et al., 2007; Antony et al., 2010; Hu et al., 2014; Zhou et al., 2015; Peng et al., 2019). The two dTALes, dT1 and dT2, were designed to specifically target a TATA box region and an upstream region of the TATA box in the promoter of *gl3*, respectively. In addition, A “T” preceding each dTALe binding element was required (Moscou and Bogdanove, 2009). The Golden Gate TALEN assembly protocol was followed to construct the two dTALes (Cermak et al., 2011). Briefly, the kit (Golden Gate TALEN and TAL Effector Kit 2.0) consisting of 86 library vectors was ordered from Addgene (www.addgene.org). To assemble the dTALe harboring 16 repeats, first 10-repeat TAL array and second 5-repeat TAL array were constructed into the destination vectors pFUS_A and pFUS_B5, respectively. The resultant vectors, the last-repeat plasmid, and the destination vector pTAL1 were digested with Esp3I restriction enzyme (Thermo Fisher Scientific, USA) and ligated with T4 Ligase (New England Biolabs, USA) to fuse all TAL repeat arrays into the pTAL1 destination vector. The dTALes were then cloned into the broad host-range vector pHM1 and transformed into Xv1601 strain by electroporation using Bio-rad Micropulser (Peng et al., 2019).

### Bacterial culture and inoculation

Xv1601 bacteria were grown on tryptone sucrose agar medium at 28°C (Peng et al., 2016). The bacterial inoculum was prepared with the OD_600_ range from 0.2 to 0.3 in the PBS buffer for plant inoculation. The second leaf of 14-day seedlings of the inbred line A188 was inoculated with the needleless syringe infiltration method. Approximately six centimeters of the second leaf from 2 cm away from the tip to the leaf base was filled with bacterial solution.

### Quantitative RT-PCR for quantifying *gl3* expression at multiple seedling stages

Shoot or second-leaf samples from A188 seedlings were collected from 3, 4, 5, 8, 14 days after seed germination. RNA was extracted from sampled tissues with Qiagen RNeasy Plant Mini Kit (Qiagen, Germany) and treated with DNase (Qiagen, Germany) to remove DNA contamination. First-strand cDNA was synthesized using Thermo Scientific Verso cDNA Kit (Thermo Fisher Scientific, USA) with anchored oligo dT primers. Quantitative RT-PCR was performed with *gl3* specific primers (**Supplemental Table 3**) and iQ™ SYBR^®^ Green Supermix (BioRad, USA) and conducted on a BioRad CFX with 96-well reaction blocks under the following PCR conditions: 95°C for 3 min, followed by 40 cycles of 15s at 95°C and 30s at 55°C. The *Actin* gene with the *actin* primers (**Supplemental Table 3**) was used as a reference gene to normalize *gl3* expression levels. The mean cycle threshold values (Ct) from technical replicates were used to calculate relative gene expression. The relative *gl3* expression was determined using the formula 100×2^(Ct_actin_ - Ct_gl3_), where Ct_actin_ and Ct_gl3_ represent the Ct values of *actin* and *gl3*, respectively.

### qRT-PCR for quantification of *gl3* expression upon dTALe treatments

To examine the expression induction of two dTALes, the second leaf of 14-day seedlings were inoculated with the bacteria containing dT1, dT2, and EV. Inoculated leaf tissues except for the inoculation position were collected at 24 h post inoculation. Three plants with the same treatment were pooled in a tissue sample. The bacterium with dT1 was used to examine the expression induction at multiple time points after the inoculation of the bacterium. Three biological replicates were conducted with three plants in each group. The inoculated leaf tissues were collected at 6 h, 12 h, 24 h, and 48 h post inoculation. qRT-PCR as mentioned was employed for the quantification of *gl3* expression.

### RNA-Seq experiment and data analysis

An RNA-Seq experiment was performed to understand the *gl3* downstream gene regulation. The bacterial inoculum was prepared to the 0.2-0.30 OD_600_ range in the PBS buffer. The second leaf of 14-day old seedlings were inoculated with a needleless syringe infiltration method. Three biological replicates (R1, R2, R3) were conducted for each of three treatment groups of which bacteria separately carried dT1, dT2, EV (empty vector). Inoculated leaf tissues were collected at 24 hours post inoculation. Three plants with the same treatment were pooled in a tissue sample. As a result, nine tissue samples were collected in total. RNA were extracted from sampled tissues with Qiagen RNeasy Plant Mini Kit. Sequencing libraries were prepared and sequenced on a Novaseq 6000 Illumina platform at Novogene Inc.. Adaptor sequences and low-quality bases of raw reads were trimmed by Trimmomatic (version 0.38) (Bolger et al., 2014). Trimmed reads were aligned to the B73 reference genome (B73Ref4) using STAR (2.7.3a) (Dobin et al., 2013). Uniquely mapped reads were used for counting reads per gene. DESeq2 (version 1.26.0) was used to identify differentially expressed genes between each of the two dTALe groups (dT1 and dT2) and the EV group. Multiple tests were accounted for by the false discovery rate (FDR) method with the FDR cutoffs of 5% for dT1 and 10% for dT2 (Benjamini and Hochberg, 1995).

### Glossy phenotyping

The glossy phenotype was identified by spraying water on seedlings at the two or three leaf stage. Seedlings whose leaves were covered with small water droplets were identified as glossy mutants.

### Scanning electron microscopy (SEM)

The second leaves collected from *ems4-12ff6f* mutant and wildtype plants were used for scanning electron microscopy analysis (HITACHI, Japan) (Aharoni et al., 2004).

### Analysis of wax composition

Wax extraction and gas chromatography-mass spectrometry (GC-MS) analyses were performed according to the described methods with some modifications (Chen et al., 2017). The *ems4-12ff6f* mutant and wildtype plants were grown in the substrate of roseite and sand (1:1) at a growth chamber (25□) until the three-leaf stage. The second leaves (about 300 mg) were collected and immersed in 3 mL of chloroform for 1 min, which dissolved 15 μg nonadecanoic acid (C19) as internal standards. The solvents were transferred into new vials and evaporated under a gentle stream of nitrogen gas. The residue was derivatized with 100 μL of N-methyl-N-(trimethylsilyl) trifluoroacetamide and incubated for 1 h at 50□. These derivatized samples were then analyzed by GC-MS (Agilent gas chromatograph coupled to an Agilent 5975C quadrupole mass selective detector). The area of leaves was calculated by IMAGEJ software (http://imagej.nih.gov/ij/). The amount of leaf wax was related to unit surface area.

### Prediction of *gl3* regulated genes through deep learning

In total, 739 B73 paired-end RNA-Seq data from diverse tissues and treatments were collected from the NCBI Sequence Read Archive (SRA) database (**Supplemental Data Set 4**). Software Trimmomatic (version 0.38) (Bolger et al., 2014) was used to trim the adaptor sequence and low-quality bases of raw reads. Remaining paired-end reads were aligned to B73 reference genome (B73Ref4) (Jiao et al., 2017) using STAR (version 2.6.0) requiring concordant mapping positions of paired-end reads (Dobin et al., 2013). Raw read counts per gene were calculated by STAR and then normalized by the library sizes of RNA-Seq samples to represent gene expression.

The 2,140 pairs of TFs and their putative targeted gene in maize obtained by homologous mapping of Arabidopsis experimented verified regulatory gene pairs from the Arabidopsis Regulatory Network database (Yilmaz et al., 2011) were used as the positive training data set for deep learning. To generate a negative data set, we randomly generated 2,140 gene pairs that do not contain above positive relationships. The maize transcriptomic data of these 4,280 gene pairs used for training the convolutional neural networks (CNN) model for predictions. The input data set of these 4,280 pairs of genes were from 739 B73 RNA-seq data. Therefore, the data set contains 4,280 gene pairs and each gene has 739 features. 428 gene pairs and their expression data, which account for 10% of the whole data set, were randomly drawn and used as the validation data set. The test data set, which contains *gl3* versus 146 *gl3* downstream genes, and also *gl3* versus 594 *gl3*-unaffected genes, were extracted from the 739 RNA-Seq data set.

Besides expression data, two additional dimensions, the product and the absolute difference of each pair of two genes, *gl3* and a putative target gene, were calculated and added as additional features. We employed Keras and TensorFlow libraries to develop the convolutional neural networks (CNN) using R libraries. The architecture of the CNN includes three parts: the input, feature extractor and classifier. The feature extractor contains several building blocks, each containing a convolution layer and a pooling layer. A convolution layer consists of multiple filters that help identify features, and activation functions that are to convert linear input to non-linear output. A pooling layer provides the down-sampling operation to reduce the dimensions of the feature map. The classifier is made up of a flatten layer and several fully connected (FC) layers and each FC layer is followed by an activation function. The flatten layer takes the results from feature extractor process and flatten them into a single long vector that can be an input for the next FC layer, which applies the weights of input to predict the true regulatory relationships and delivers the final output of the network as represented by probability for each pair of genes for prediction. To identify a model with a high performance for the prediction, we tried multiple loss functions. The mean squared logarithmic error loss (MSLE) was selected as the loss function.

### Inference of Gl3-regulated target genes using top-down GGM algorithm

The top-down Gaussian graphical model (GGM) algorithm developed earlier (*Wei, 2019; Lin et al., 2013*) was employed to construct a multilayered gene regulatory network (ML-hGRN) mediated by GL3 in two steps, with the dT1, dT2, EV RNA-seq data being used as the input data. Briefly, in the first step, the GL3 downstream genes that responded to the *gl3* activation were identified using Fisher’s exact test and the probability-based method as we described in our publications mentioned above, and these genes were termed *gl3* responsive genes; in the second step, we further identified those that were interfered frequently by GL3 from the *gl3* responsive genes through evaluating all triple gene blocks, each consisting of *gl3*, defined as z, and two *gl3* responsive genes, defined as x and y. If GL3 significantly interfered with the two responsive genes in a triple gene block, the difference (d) between the correlation coefficient, *r_xy_* of two responsive genes in expression and the partial correlation coefficient, *r_xy/z_*, representing the correlation of two *gl3* responsive genes conditioning on *gl3* (*z*) should be significant. The null hypothesis H_0_: *d* = 0 was tested with the multivariate delta method (MacKinnon et al., 2002). If *d* is significantly different from 0, *gl3* was concluded to interfere with the two responsive genes and their regulatory relationships were recorded once. After all combined triple gene blocks were evaluated, the interference frequency between *gl3* and each responsive gene was calculated. In this study, the candidate target genes with at least one inference frequency were considered to be a gene directly regulated by *gl3*.

The Metabolomics Facility of the Institute of Genetics and Developmental Biology, Chinese Academy of Sciences

## ACKNOWLEDGEMENTS

We thank Fengxia Zhang from the Metabolomics Facility of the Institute of Genetics and Developmental Biology at Chinese Academy of Sciences for wax composition analysis, Wei Wang from Eduard Akhunov laboratory and Melinda Dalby from Barbara Valent laboratory at Kansas State University for helping on confocal microscope experiments. We thank funding support from the US National Science Foundation (awards no. 1741090), the USDA’s National Institute of Food and Agriculture (award no. 2018-67013-28511), and the Agricultural Science and Technology Innovation Program of Chinese Academy of Agricultural Sciences. This is the contribution number 21-196-J from the Kansas Agricultural Experiment Station.

## AUTHOR CONTRIBUTIONS

J.Z., M.C., S.P., H.W., F.F.W, and S.L. conceived and designed experiments. M.Z., Z.P., Y.Q., B.T., Y.C., G.L., H.Z., K.L., H.T., Y.L., and J.Z. performed experiments and collected data. M.Z., Z.P., Y.Q., L.Z., C.H., H.W., and S. L. analyzed data. M.Z., Z.P., Y.Q., L.Z., Y.L., M.C., S.P., J.Z., H.W., F.F.W., and S.L. wrote the manuscript with comments from other authors.

## DATA AVAILABILITY

Raw dTALe RNA-Seq data are available at NCBI SRA under the project of PRJNA692729.

## SUPPLEMENTARY INFORMATION

Supplemental Figures: Supplemental Figures 1-5

Supplemental Tables: Supplemental Tables 1-3

Supplemental Data Set 1: Detailed information of DEGs

Supplemental Data Set 2: Deep learning classification for GL3 downstream genes and dTALe unaffected genes

Supplemental Data Set 3: Analyzing result from the top-down GGM of gl3 downstream genes

Supplemental Data Set 4: List of 739 B73 RNA-Seq data accessions used for deep learning

## Parsed Citations

Aharoni, A, Dixit, S., Jetter, R., Thoenes, E., van Arkel, G., and Pereira, A. (2004). The SHINE clade of AP2 domain transcription factors activates wax biosynthesis, alters cuticle properties, and confers drought tolerance when overexpressed in Arabidopsis. Plant Cell 16: 2463–2480.

Antony, G., Zhou, J., Huang, S., Li, T., Liu, B., White, F., and Yang, B. (2010). Rice xa13 recessive resistance to bacterial blight is defeated by induction of the disease susceptibility gene Os-11N3. Plant Cell 22: 3864–3876.

Bartlett, A, O’Malley, R.C., Huang, S.-S.C., Galli, M., Nery, J.R., Gallavotti, A, and Ecker, J.R. (2017). Mapping genome-wide transcription-factor binding sites using DAP-seq. Nat. Protoc. 12: 1659–1672.

Benjamini, Y. and Hochberg, Y. (1995). Controlling the false discovery rate: a practical and powerful approach to multiple testing. J. R. Stat. Soc. Series B Stat. Methodol. 57: 289–300.

Block, A, Li, G., Fu, Z.Q., and Alfano, J.R. (2008). Phytopathogen type III effector weaponry and their plant targets. Curr. Opin. Plant Biol. 11: 396–403.

Boch, J., Scholze, H., Schornack, S., Landgraf, A, Hahn, S., Kay, S., Lahaye, T., Nickstadt, A, and Bonas, U. (2009). Breaking the code of DNA binding specificity of TAL-type III effectors. Science 326: 1509–1512.

Bolger, AM., Lohse, M., and Usadel, B. (2014). Trimmomatic: a flexible trimmer for Illumina sequence data. Bioinformatics 30: 2114–2120.

Büttner, D. and Bonas, U. (2010). Regulation and secretion of Xanthomonas virulence factors. FEMS Microbiol. Rev. 34: 107–133.

Cermak, T., Doyle, E.L., Christian, M., Wàng, L., Zhang, Y., Schmidt, C., Baller, J.A, Somia, N.V., Bogdanove, AJ., and Voytas, D.F. (2011). Efficient design and assembly of custom TALEN and other TAL effector-based constructs for DNA targeting. Nucleic Acids Res. 39: e82.

Chen, X. et al. (2017). IRREGULAR POLLEN EXINE1 Is a novel factor in anther cuticle and pollen exine formation. Plant Physiol. 173: 307–325.

Costa, T.R.D., Felisberto-Rodrigues, C., Meir, A, Prevost, M.S., Redzej, A, Trokter, M., and Waksman, G. (2015). Secretion systems in Gram-negative bacteria: structural and mechanistic insights. Nat. Rev. Microbiol. 13: 343–359.

Cunningham, F.J., Goh, N.S., Demirer, G.S., Matos, J.L., and Landry, M.P. (2018). Nanoparticle-mediated delivery towards advancing plant genetic engineering. Trends Biotechnol. 36: 882–897.

Demirer, G.S., Zhang, H., Goh, N.S., Pinals, R.L., Chang, R., and Landry, M.P. (2020). Carbon nanocarriers deliver siRNA to intact plant cells for efficient gene knockdown. Sci Adv 6: eaaz0495.

Deslandes, L. and Rivas, S. (2012). Catch me if you can: bacterial effectors and plant targets. Trends Plant Sci. 17: 644–655.

Dobin, A, Davis, C.A., Schlesinger, F., Drenkow, J., Zaleski, C., Jha, S., Batut, P., Chaisson, M., and Gingeras, T.R. (2013). STAR: ultrafast universal RNA-seq aligner. Bioinformatics 29: 15–21.

Fehling, E. and Mukherjee, K.D. (1991). Acyl-CoA elongase from a higher plant (Lunaria annua): metabolic intermediates of very-long-chain acyl-CoA products and substrate specificity. Biochim. Biophys. Acta 1082: 239–246.

Gleba, Y.Y., Tusé, D., and Giritch, A (2014). Plant viral vectors for delivery by Agrobacterium. Curr. Top. Microbiol. Immunol. 375: 155–192.

Green, E.R. and Mecsas, J. (2016). Bacterial secretion systems: an overview. Microbiol. Spectr. 4: 1–32.

Hansen, J.D., Pyee, J., Xia, Y., Wen, T.J., Robertson, D.S., Kolattukudy, P.E., Nikolau, B.J., and Schnable, P.S. (1997). The glossy1 Locus of maize and an epidermis-specific cDNA from Kleinia odora Define a class of receptor-like proteins required for the normal accumulation of cuticular waxes. Plant Physiol. 113: 1091–1100.

Hu, Y., Zhang, J., Jia, H., Sosso, D., Li, T., Frommer, W.B., Yang, B., White, F.F., Wang, N., and Jones, J.B. (2014). Lateral organ boundaries 1 is a disease susceptibility gene for citrus bacterial canker disease. Proc. Nat. Acad. Sci. U. S. A 111: E521–E529.

Jiao, Y. et al. (2017). Improved maize reference genome with single-molecule technologies. Nature 546: 524–527.

Joung, J.K. and Sander, J.D. (2013). TALENs: a widely applicable technology for targeted genome editing. Nat. Rev. Mol. Cell Biol. 14: 49–55.

Kay, S., Hahn, S., Marois, E., Hause, G., and Bonas, U. (2007). A bacterial effector acts as a plant transcription factor and induces a cell size regulator. Science 318: 648–651.

Khang, C.H., Berruyer, R., Giraldo, M.C., Kankanala, P., Park, S.-Y., Czymmek, K., Kang, S., and Valent, B. (2010). Translocation of Magnaporthe oryzae effectors into rice cells and their subsequent cell-to-cell movement. Plant Cell 22: 1388–1403.

Kunst, L. and Samuels, A.L. (2003). Biosynthesis and secretion of plant cuticular wax. Prog. Lipid Res. 42: 51–80.

Lai, X., Stigliani, A, Vachon, G., Carles, C., Smaczniak, C., Zubieta, C., Kaufmann, K., and Parcy, F. (2019). Building transcription factor binding site models to understand gene regulation in plants. Mol. Plant 12: 743–763.

Lee, S.B. and Suh, M.C. (2013). Recent advances in cuticular wax biosynthesis and its regulation in Arabidopsis. Mol. Plant 6: 246–249.

Li, L., Du, Y., He, C., Dietrich, C.R., Li, J., Ma, X., Wang, R., Liu, Q., Liu, S., Wang, G., and Others (2019). Maize glossy6 is involved in cuticular wax deposition and drought tolerance. J. Exp. Bot. 70: 3089–3099.

Li, L., Li, D., Liu, S., Ma, X., Dietrich, C.R., Hu, H.-C., Zhang, G., Liu, Z., Zheng, J., Wang, G., and Schnable, P.S. (2013a). The maize glossy13 gene, cloned via BSR-Seq and Seq-walking encodes a putative ABC transporter required for the normal accumulation of epicuticular waxes. PLoS One 8: e82333.

Lin, Y.C., Li, W., Sun, Y.H., Kumari, S., Wei, H., and Li, Q. (2013). SND1 transcription factor–directed quantitative functional hierarchical genetic regulatory network in wood formation in Populus trichocarpa. The Plant 25: 4324–4341.

Li, T., Huang, S., Zhou, J., and Yang, B. (2013b). Designer TAL effectors induce disease susceptibility and resistance to Xanthomonas oryzae pv. oryzae in rice. Mol. Plant 6: 781–789.

Liu, S., Dietrich, C.R., and Schnable, P.S. (2009). DLA-based strategies for cloning insertion mutants: cloning the gl4 locus of maize using Mu transposon tagged alleles. Genetics 183: 1215–1225.

Liu, S., Yeh, C.-T., Tang, H.M., Nettleton, D., and Schnable, P.S. (2012). Gene mapping via bulked segregant RNA-Seq (BSR-Seq). PLoS One 7: e36406.

Lu, X. et al. (2018). Gene-indexed mutations in maize. Mol. Plant 11: 496–504.

MacKinnon, D.P., Lockwood, C.M., Hoffman, J.M., West, S.G., and Sheets, V. (2002). A comparison of methods to test mediation and other intervening variable effects. Psychol Methods 7: 83–104.

Minsavage, G.V. (1990). Gene-for-Gene relationships specifying disease resistance in Xanthomonas campestris pv. vesicatoria - Pepper Interactions. Mol. Plant Microbe Interact. 3: 41–47.

Moose, S.P. and Sisco, P.H. (1996). Glossy15, an APETALA2-like gene from maize that regulates leaf epidermal cell identity. Genes Dev. 10: 3018–3027.

Moscou, M.J. and Bogdanove, AJ. (2009). A simple cipher governs DNA recognition by TAL effectors. Science 326: 1501.

Peng, Z., Hu, Y., Xie, J., Potnis, N., Akhunova, A., Jones, J., Liu, Z., White, F.F., and Liu, S. (2016). Long read and single molecule DNA sequencing simplifies genome assembly and TAL effector gene analysis of Xanthomonas translucens. BMC Genomics 17: 21.

Peng, Z., Hu, Y., Zhang, J., Huguet-Tapia, J.C., Block, A.K., Park, S., Sapkota, S., Liu, Z., Liu, S., and White, F.F. (2019). Xanthomonas translucens commandeers the host rate-limiting step in ABA biosynthesis for disease susceptibility. Proc. Natl. Acad. Sci. U. S. A 116: 20938–20946.

Perez-Quintero, AL. et al. (2020). Genomic acquisitions in emerging populations of Xanthomonas vasicola pv. vasculorum Infecting corn in the United States and Argentina. Phytopathology 110: 1161–1173.

Sugio, A, Yang, B., Zhu, T., and White, F.F. (2007). Two type III effector genes of Xanthomonas oryzae pv. oryzae control the induction of the host genes OsTFIIA 1 and OsTFX1 during bacterial blight of rice. Proceedings of the National Academy of Sciences 104: 10720–10725.

Tacke, E., Korfhage, C., Michel, D., Maddaloni, M., Motto, M., Lanzini, S., Salamini, F., and Döring, H.P. (1995). Transposon tagging of the maize Glossy2 locus with the transposable element En/Spm. Plant J. 8: 907–917.

Tang, J., Yang, X., Xiao, C., Li, J., Chen, Y., Li, R., Li, S., Lü, S., and Hu, H. (2020). GDSL lipase Occluded Stomatal Pore 1 is required for wax biosynthesis and stomatal cuticular ledge formation. New Phytol. 228: 1880–1896.

Tran, T.T. et al. (2018). Functional analysis of African Xanthomonas oryzae pv. oryzae TALomes reveals a new susceptibility gene in bacterial leaf blight of rice. PLoS Pathog. 14: e1007092.

Van den Ackerveken, G., Marois, E., and Bonas, U. (1996). Recognition of the bacterial avirulence protein AvrBs3 occurs inside the host plant cell. Cell 87: 1307–1316.

Wei, H. (2019). Construction of a hierarchical gene regulatory network centered around a transcription factor. Brief. Bioinform 20: 1021–1031.

Wei, M. et al. (2020). PuHox52-mediated hierarchical multilayered gene regulatory network promotes adventitious root formation in Populus ussuriensis. New Phytol. 228: 1369–1385.

White, F.F., Potnis, N., Jones, J.B., and Koebnik, R. (2009). The type III effectors of Xanthomonas. Mol. Plant Pathol. 10: 749–766.

Xu, X., Dietrich, C.R., Delledonne, M., Xia, Y., Wen, T.J., Robertson, D.S., Nikolau, B.J., and Schnable, P.S. (1997). Sequence analysis of the cloned glossy8 gene of maize suggests that it may code for a [beta]-ketoacyl reductase required for the biosynthesis of cuticular waxes. Plant Physiol. 115: 501–510.

Yang, B., Zhu, W., Johnson, L.B., and White, F.F. (2000). The virulence factor AvrXa7 of Xanthomonas oryzae pv. oryzae is a type III secretion pathway-dependent nuclear-localized double-stranded DNA-binding protein. Proc. Natl. Acad. Sci. U. S. A 97: 9807–9812.

Yilmaz, A, Mejia-Guerra, M.K., Kurz, K., Liang, X., Welch, L., and Grotewold, E. (2011). AGRIS: the Arabidopsis Gene Regulatory Information Server, an update. Nucleic Acids Res. 39: D1118–1122.

Zheng, J., He, C., Qin, Y., Lin, G., Park, W.D., Sun, M., Li, J., Lu, X., Zhang, C., Yeh, C.-T., and Others (2019). Co-expression analysis aids in the identification of genes in the cuticular wax pathway in maize. Plant J. 97: 530–542.

Zhou, J., Peng, Z., Long, J., Sosso, D., Liu, B., Eom, J.-S., Huang, S., Liu, S., Vera Cruz, C., Frommer, W.B., White, F.F., and Yang, B. (2015). Gene targeting by the TAL effector PthXo2 reveals cryptic resistance gene for bacterial blight of rice. Plant J. 82: 632–643.

Zhu, W., Yang, B., Chittoor, J.M., Johnson, L.B., and White, F.F. (1998). AvrXa10 contains an acidic transcriptional activation domain in the functionally conserved C terminus. Mol. Plant. Microbe. Interact. 11: 824–832.

